# Co-opting MBNL-dependent alternative splicing cassette exons to control gene therapy in myotonic dystrophy

**DOI:** 10.1101/2025.01.23.634565

**Authors:** ST Carrell, EM Carrell, RC Giovenco, BL Davidson

## Abstract

Myotonic dystrophy type 1 (DM1) is a multisystemic genetic disorder caused by a CTG repeat expansion that accumulate as toxic CUG repeat RNA. Functional sequestration of muscleblind-like (MBNL) proteins by CUG repeat RNA leads to deleterious, yet predictable changes in alternative splicing in DM. Genetic medicines for DM that reduce CUG repeat RNA or increase MBNL are advancing, but application of a viral-based approach must contend with high phenotypic and molecular variability. To address this, we have repurposed well-described cassette exons that are excluded by MBNL action to tune translational output of a therapeutic protein. We show that these splicing events can be taken out of genetic context and respond to changes in MBNL concentration or accumulation of toxic CUG repeat RNA, can deliver therapeutic MBNL1 protein to improve skeletal muscle myotonia or prevent cardiac toxicity associated with MBNL1 overexpression in mice, and distinguish DM patient-derived skeletal muscle myotubes from isogenic controls. Further work to fine-tune events to specific tissue targets and therapeutic cargos is needed, but these events can increase the therapeutic window for viral-based approaches for DM1.

## Introduction

Genetic therapies have become the standard-of-care for multiple inherited neuromuscular disorders, including spinal muscular atrophy^1^, Duchenne muscular dystrophy^2^ and transthyretin-related amyloidosis neuropathy^3^. Of these, viral-based approaches have the ability to efficiently deliver throughout the body and have a prolonged action when targeting post-mitotic cells. These advantages come with drawbacks, including immunogenicity that inhibits redosing regardless of efficacy and inability to remove or silence the therapy if toxic.

To improve efficacy and limit toxicity, efforts have been focused on developing novel viral capsids that reduce delivery to non-target tissues^4^, cargo elements that direct tissue-specific expression, or cargo elements that rely on post-transcriptional RNA regulation, including elements that lead to alternative splicing in response to small molecules^5^ or alterations in splicing factor function^6,7^.

Myotonic dystrophy type 1 (DM1) is an autosomal dominant genetic disorder that results in progressive muscular weakness and atrophy, cardiac conduction disease, and a multitude of other variable systemic effects. DM1 is caused by expansion of a non-coding CTG repeat in *DMPK* that is transcribed and accumulates as toxic CUG repeat RNA. The CUG RNA exerts its toxicity through binding and sequestration of RNA-binding proteins of the muscleblind-like (MBNL) family, which are involved in pre-mRNA processing^8,9^, including alternative splicing. Alternative splicing changes that occur due to MBNL loss-of-function in DM1 have been shown to cause specific disease phenotypes^10,11^ and can serve as quantitative biomarkers of disease in DM1 skeletal muscle^12^.

Genetic medicines to reduce toxic RNA accumulation are in early phase clinical trials for DM1, and, similar to other inherited neuromuscular conditions, a long-acting viral-based gene therapy may offer additional benefit. The application of gene therapy to DM1, however, poses several hurdles, as DM1 is a multisystemic disease with high variability^13^. To address this hurdle, we have developed control elements that can post-transcriptionally control a gene therapy for DM1 by responding to MBNL protein activity (DMX^on^). DMX^on^ elements are derived from well-documented MBNL-dependent splicing events^12,14^, and through addition of a strong Kozak sequence, can be placed upstream of any protein-coding transgene to control its translation. Here, we provide evidence that alternative splicing cassettes retain partial activity once removed from their natural context, can be tuned to alter their responses *in vitro* and *in vivo*, and can be used to adjust protein output to limit toxicity or promote therapeutic efficacy in mouse models and iPSC-derived skeletal myotubes of DM1.

## Results

### Identification and *in vitro* evaluation of DMX^on^ candidates

To identify DMX^on^ candidates we considered MBNL-dependent alternative splicing events characterized as biomarkers of disease severity in skeletal muscle^11,12,14–17^. These events were shown to correlate with weakness in tibialis anterior (TA) muscles of patients with DM1^12^ and have been evaluated in both MBNL loss-of-function and toxic RNA gain-of-function mouse models^14^. To narrow our selection, we chose splicing events based on 4 criteria: (1) is a single, short alternate exon (i.e. could not be a mutual or more complex event), (2) is excluded by MBNL, (3) has a relatively large change in percent-spliced-in (PSI) across the spectrum of disease severity (>30% *Δ*PSI), and (4) is consistent across different skeletal muscles^14^. Within these criteria our list was narrowed to 7 events: *LDB3* exon 8, *DCTN4* exon 6, *MPDZ* exon 27, *NFIX* exon 7, *MBNL1* exon 5, *JAG2* exon 10, and *MAP3K4* exon 17. Of these 7, we chose 3 events to evaluate as minigenes *in vitro. LDB3* exon 8 was chosen as it had the largest *Δ*PSI, *MBNL1* exon 5 was chosen for its low EC50 and steep slope in response to [MBNL], and *NFIX* exon 7 was chosen for its high EC50 and relatively shallow slope of response to [MBNL]. *JAG2* and *MAP3K4* had similar range and slope of response to *MBNL1* exon 5, albeit with larger alternate exons. *DCTN4* and *MPDZ* behaved similar to *NFIX* exon 7, but the small size of *DCTN4* exon 6 (21 nt) raised concern about the effects of Kozak sequence insertion, as the alternate exon is known to contribute to splicing response in MBNL-dependent exclusion events^8^.

Alternative splicing is a co-transcriptional event that occurs during transcript elongation. Splicing efficiency of an intron depends on splicing sequences (5’ - and 3’ - splice sites, branch point, exonic and intronic splicing enhancers) and other mRNA maturation steps, thus it is context dependent^18^. To first ascertain MBNL dose-response of splicing cassettes outside their natural pre-mRNA context, we generated minigenes from human gDNA sequence of *NFIX* exon 6-8, *MBNL1* exon 4-6, and *LDB3* exon 7-9. The minigenes were modified to include a strong Kozak sequence within the alternate exon, in frame with the natural coding sequence, and cloned upstream an eGFP reporter cDNA lacking a start codon (Figure 1A). A tetracycline-inducible HA-MBNL1 HEK 293 cell line^17^ (kind gift of Andrew Berglund) was used to titrate [MBNL1] and determine minigene responses (Supplemental Figure 1A). Similar to prior observations, NFIX had the highest EC50 of the three events, and LDB3 had the largest PSI range (Figure 1B) across a range of [MBNL1]. Unexpectedly, gel analysis of LDB3 had an extra, larger amplicon band suggesting downstream intron retention; therefore, this event was not evaluated further (Supplemental Figure 1B).

**Fig 1.**
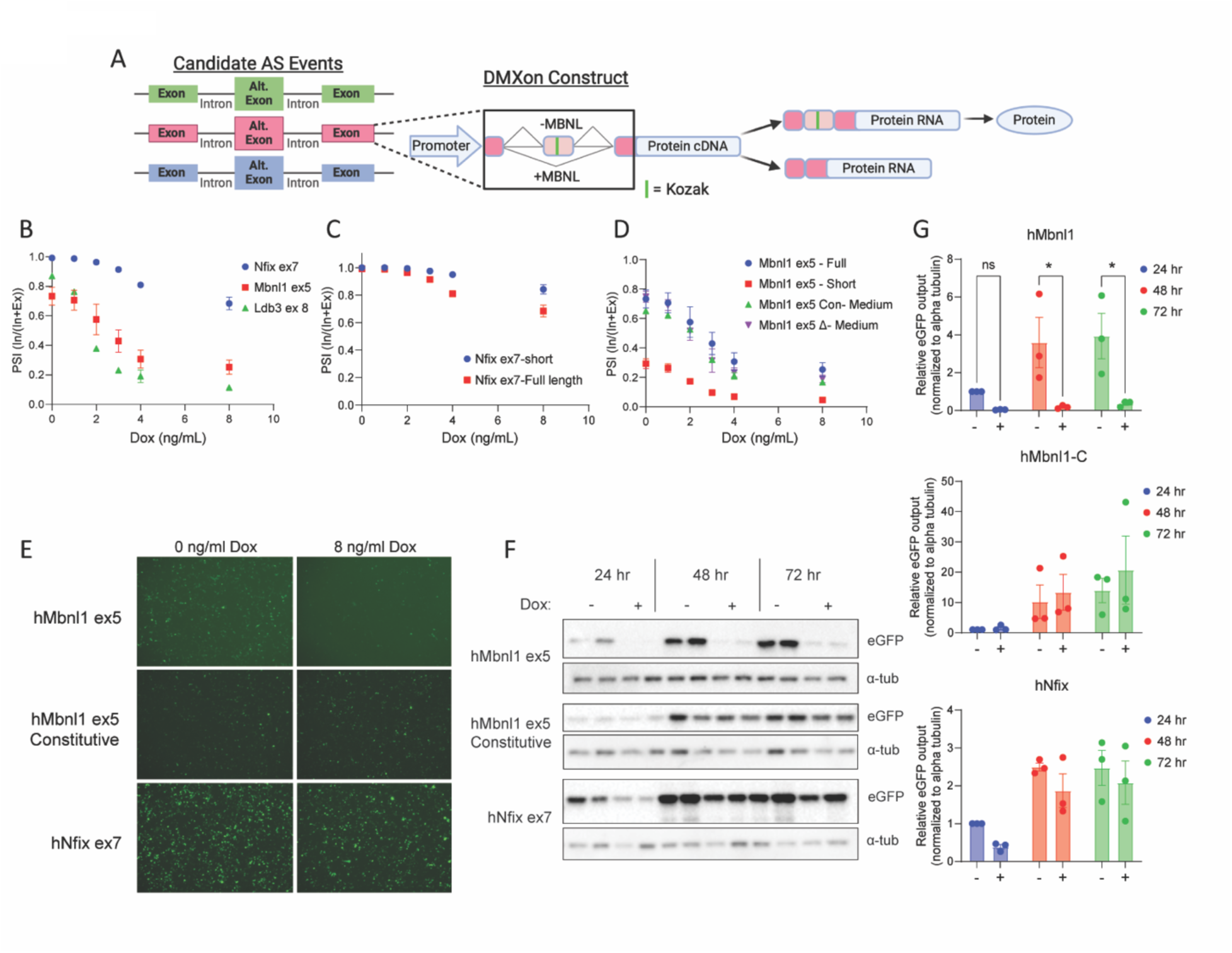
DMX^on^ Design and *in vitro* testing. (A) Diagram of DMX^on^ splicing cassette design. Candidate MBNL-dependent alternative splicing events removed from natural context, strong Kozak sequence inserted in alternate exon (green bar). (B) Quantification of percent spliced in (PSI) for full-length DMX^on^ events (Nfix exon 7, Mbnl1 exon 5, LDB3 exon 8) across a range of doxycycline doses in tet-on HA-MBNL1 HEK293 cells. (C) As in (B), Nfix exon 7 with intronic truncations. (D) As in (B), Mbnl1 exon 5 with intronic truncations and mutation of Mbnl1 exon 4 5’ splice site to remove start codon (*Δ*). (E) Representative images of eGFP fluorescence 24 hours following transfection in 0 ng/mL or 8 ng/mL doxycycline. (F) Representative images of Western blot for eGFP protein across a 72-hour time course. *α*-tubulin serves as loading control. (G) Quantification of Western blot, as shown in (F).

Although we selected for events that contained short cassette exons, the introns of the full-length minigenes were too large for the genomic packaging limit of AAV (∼4.7 kb). MBNL suppression events, such as those used in our cassettes, are modulated by MBNL binding within the alternative exon itself, the ∼140 bp at the 3’ end of the upstream intron and ∼100 bp at the 3’ end of the downstream intron^19^. Using this information and a map of putative MBNL binding sites for each event (https://rbpmap.technion.ac.il/), we truncated intronic regions to develop short variants of both minigene reporters. The shortened variants of both reporters showed a reduced range of PSI across the [MBNL1] curve (Figure 1C-D). To retain the dynamic range observed in the full-length MBNL1 minigene, a third variant that reconstituted a portion of the upstream intron 4, MBNL1-medium, was generated. Further, the original MBNL1 full-length minigene contained a methionine initiation codon flanking the 5’-splice site at the boundary of exon/intron 4. This was mutated to generate the MBNL1-medium *Δ* (Figure 1D), which retains the *in vitro* splicing response of MBNL1-medium and is termed hMbnl1 going forward. The minigene retaining this start codon is termed hMBNL1-medium constitutive (hMbnl1-C), and serves as a splicing-independent positive control.

Next, we assessed whether observed changes in PSI lead to changes in reporter protein production. Fluorescent imaging of eGFP intensity at 24 hours and Western blot for eGFP protein across a 72-hour time course were performed at no (0 ng/mL dox) or high (8 ng/mL dox) [MBNL1]. Consistent with PSI measurements, high [MBNL1] suppressed eGFP production in hMBNL1, but not in the hNfix or hMBNL1-C minigenes (Figure 1E-G).

### *In vivo* DMX^on^ responds to expanded repeat RNA accumulation and shows negative feedback and improvement of myotonia when expressing therapeutic MBNL1

To assess the ability of DMX^on^ to control AAV transgene output *in vivo*, we packaged NFIX and MBNL1 minigenes that express eGFP or therapeutic MBNL1 protein and were driven by the skeletal and cardiac muscle-specific promoter, CK8. MBNL1 overexpression was chosen as a therapeutic strategy to test DMX^on^ because it was previously shown to reduce muscle myotonia in human skeletal actin long-repeat (HSA^LR^) mice following AAV delivery by intramuscular (IM) injection^20^, and for the purposes of testing a transgene control switch, it is known to cause cardiotoxicity when delivered systemically to wild-type (WT) mice^21^.

DMX^on^-controlled vectors were packaged into AAV9 and delivered by intramuscular injection (1E9 vg) into TA muscles of WT or HSA^LR^ mice. The HSA^LR^ mouse is a commonly used skeletal muscle model of myotonic dystrophy that expresses a hACTA1 transgene containing ∼220 CTG repeats in the 3’ untranslated region^22^. Transgene expression leads to CUG repeat RNA production, MBNL sequestration, alternative splicing changes, and hind limb myotonia^11^.

Initial evaluation of eGFP-expressing vectors in TA muscles showed that the hMbnl1 minigene distinguished between WT and HSA^LR^ mice and that transgene-derived MBNL1 expression reduced cassette exon inclusion (Figure 2A). In contrast, the hNfix cassette was less sensitive to differences in [MBNL], showing nearly 100% exon inclusion in both mouse strains. Further, transgene-derived expression of MBNL1 was insufficient to reduce exon inclusion (Figure 2B). These differences are consistent with observations of self-inhibitory feedback observed over a 72-hour time course *in vitro* (Supplemental Figure 2).

**Fig 2.**
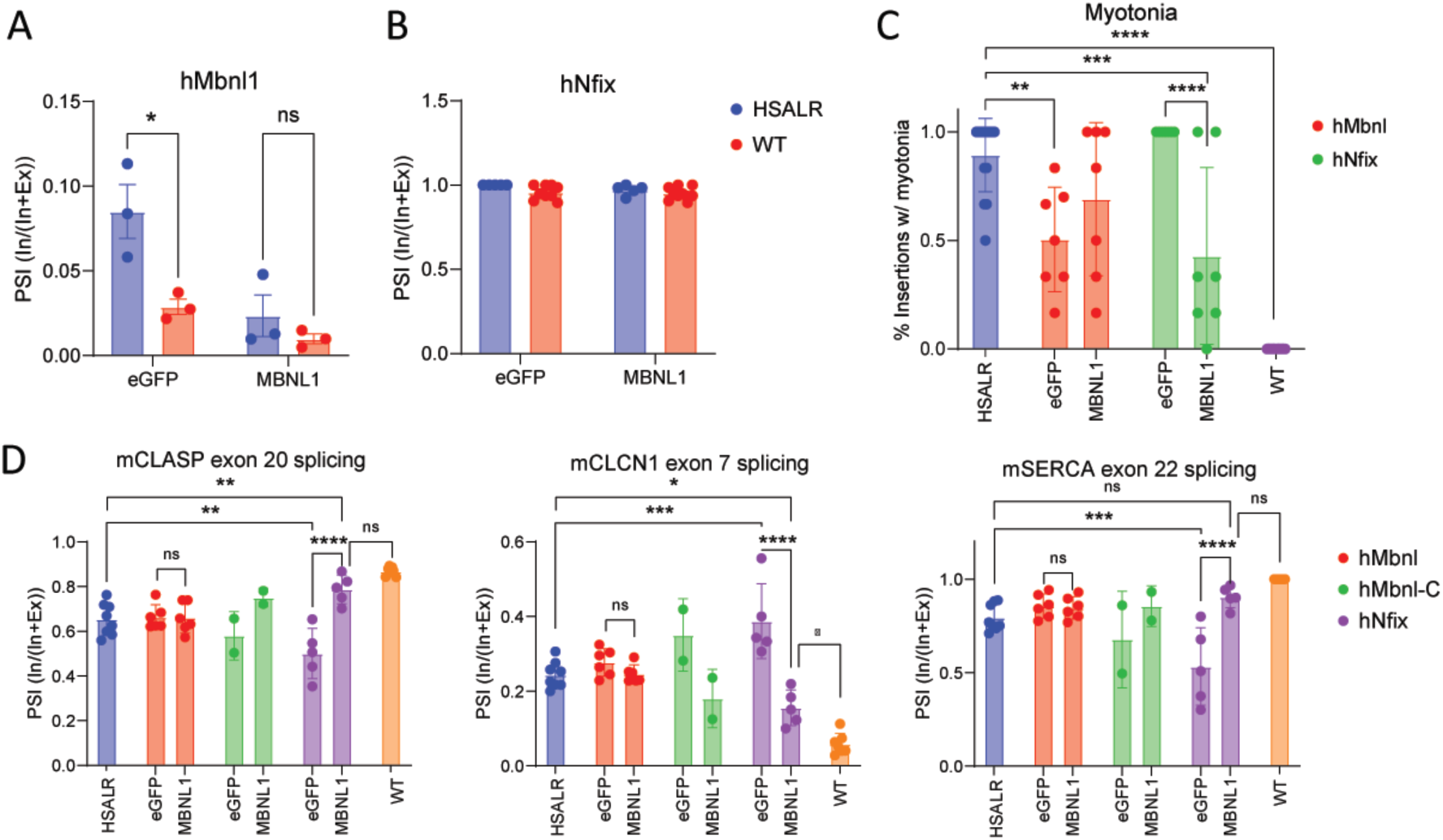
AAV9-delivered DMX^on^ cassettes respond differently to expanded repeats *in vivo*. (A) Quantification of PSI of DMX^on^ hMbnl1 exon 5 and hNfix exon 7 minigenes by ddPCR in tibialis anterior (TA) muscles of WT or HSA^LR^ mice when expressing eGFP reporter or MBNL1 therapeutic protein (n = 5-6/group). (B) Quantification of myotonia by needle EMG as percentage of insertions leading to a myotonic discharge (n = 7-14/group). (C) Quantification of endogenous MBNL-dependent splicing events [Clasp1 exon 20, Clcn1 exon 7, and Serca exon 22] to assess therapeutic effect at 12 weeks post-injection (n=2-8). Individual pairwise comparisons were performed with Student’s T-test for B-C. For D, one-way ANOVA followed by Sidak’s multiple comparisons was performed. * p<0.05, ** p<0.01,*** p<0.005, **** p<0.0001.

The therapeutic impact of MBNL1 overexpression was assessed by measurements of myotonia by needle electromyography (EMG) and evaluation of known endogenous splicing events, *Clasp1* exon 20, *Clcn1* exon 7, and *Serca* exon 22. At 12-weeks post-injection a therapeutic effect was observed only with hNfix.MBNL1 vectors (Figure 2C,D). Interestingly, hNfix.eGFP treated animals had more severe myotonia and mis-splicing of endogenous events, and mice treated with hMbnl1.eGFP seemed to have a milder phenotype.

### DMX^on^ switches can be modified to tune their response

Because hNfix did not respond to alterations in functional [MBNL] *in vivo* and inclusion of hMbnl1 was too low to lead to phenotypic improvement *in vivo,* we sought to adjust the responsivity of both DMX^on^ switches.

For the hMbnl1 construct we mutated two locations of a known intronic splicing silencer region in exon 4 (Figure 3A), previously shown to increase PSI across a range of MBNL1 concentrations in tetracycline-inducible HA-MBNL1 HEK 293 cells^17^. Individual modifications 3M, a two G>C nucleotide substitution, del 2, an eighteen bp deletion, or both mutations combined (3M + del 2), increased PSI across a range of doxycycline doses (Figure 3B). The combination (3M + del 2) led to use of an alternate splice acceptor (Supplemental Figure 3); therefore, this variant was not pursued further. Del 2 retained the dynamic range of the original hMbnl1 construct as measured by PSI and protein output *in vitro* (Figure 3B,C) and was used as DMX^on^ hMbnl1-version 2 (hMbnl1V2).

**Fig 3.**
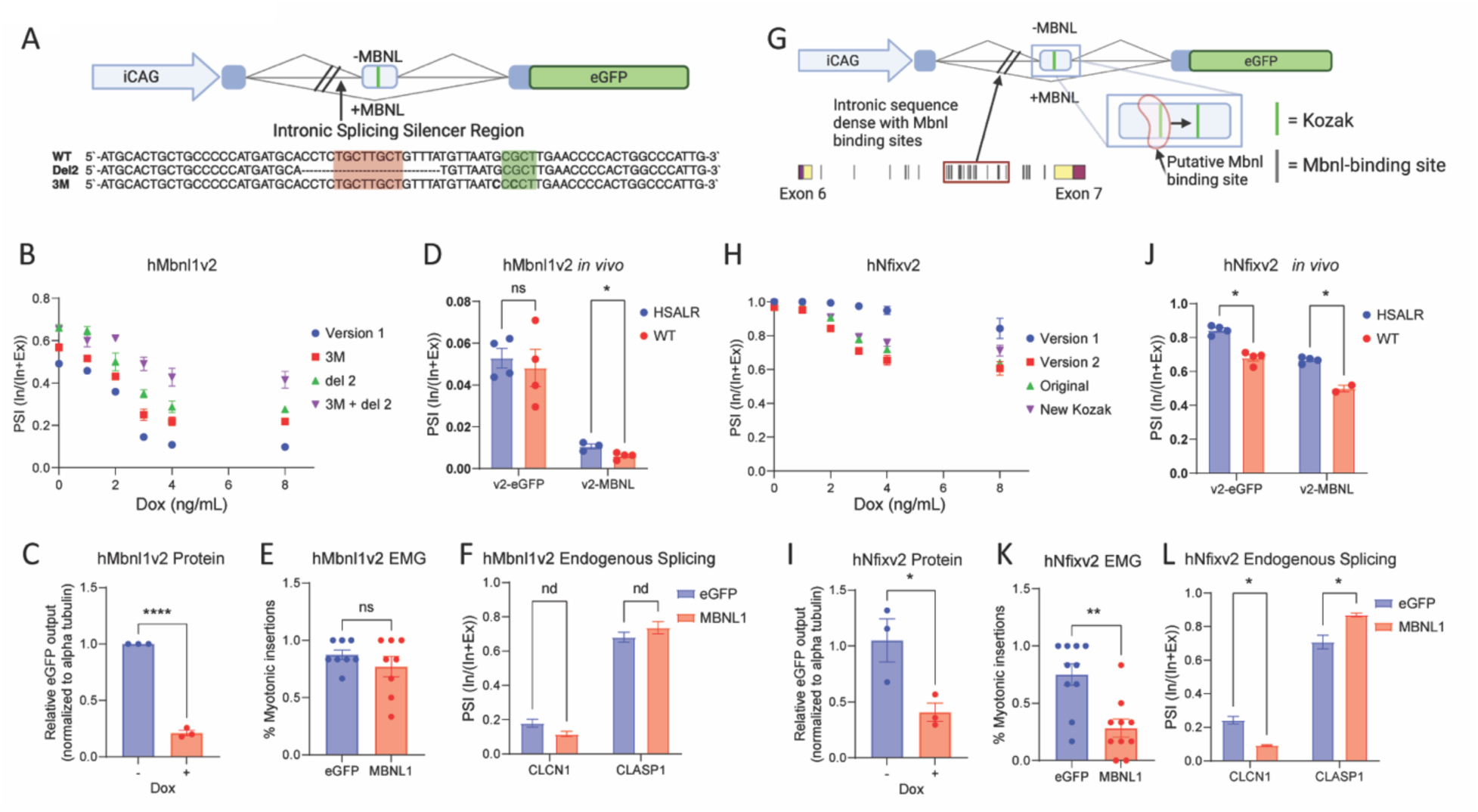
Changes to coding and non-coding regions of DMX^on^ minigenes alter their response to [MBNL]. (A) Diagram of hMbnl1 exon 5 minigene with del2 and 3M mutations in intronic splicing silencer (adapted from Wagner, et al 2016). (B) Quantification of PSI of hMbnl1 mutation variants across a range of doxycycline (n = 3 per treatment/group), and (C) eGFP protein output from iCAG.hMbnl1V2.eGFP minigene with 0 ng/mL or 8 ng/mL doxycycline conditions after 72 hours (n = 3 per treatment) in HA-MBNL1 HEK293 cells. (D) PSI of CK8.hMbnl1V2 cassettes at 12 weeks following IM injection in TA muscles of WT or HSA^LR^ mice (n = 4 muscles per treatment/genotype). (E) Myotonia measured by needle EMG 12 weeks post-IM injection of hMbnl1V2.eGFP or -.MBNL1 (n = 8 per treatment). (F) Quantification of endogenous MBNL-dependent splicing events [Clasp1 exon 20, Clcn1 exon 7] at 12 weeks following IM injection of hMbnl1V2.eGFP or -.MBNL1 (n = 3/group). (G) Diagram of hNfix exon 7 minigene with alterations to the Kozak sequence location within exon 7 and addition of sequence in intron 6. (H) Quantification of PSI of hNfix variants, compared to original full-length minigene, across a range of doxycycline (n = 3 per treatment/group). (I) Quantification of eGFP protein by WB of iCAG.hNfixV2.eGFP minigene, as in (C). (J) As in (D), measurement of PSI of CK8.hNfixV2 vectors following IM injection (n = 4 muscles per vector/genotype). (K) As in (E), with hNfixV2.eGFP or -.MBNL1 (n = 10 per treatment). (L) As in (F), with hNfix1v2.eGFP or -.MBNL1 (n = 3 / group).

Next, modifications to hNfix included moving the Kozak sequence in exon 7 downstream of a putative MBNL1 binding site and re-introducing ∼1kb of intron 6 sequence with a high density of putative MBNL1 binding sites (hNfixV2; Figure 3G). These changes led to a significant reduction in PSI and reduced eGFP protein output *in vitro* in tetracycline-inducible HA-MBNL1 HEK 293 cells (Figure 3H-I).

Contrary to the *in vitro* testing, when direct IM injections were performed of hMbnl1V2, PSI was similar to the first hMbnl1 minigene but no longer distinguished between WT and HSA^LR^ TA muscles (Figure 3D). hMbnl1V2 still did not significantly improve muscle myotonia (Figure 3E) nor *Clasp1* exon 20 splicing, and minimally altered endogenous *Clcn1* exon 7 splicing in both WT and HSA^LR^ (Figure 3F, Supplemental Figure 4A). To examine whether this reduction in hMbnl1V2 splicing response *in vivo* may be due to the CK8 promoter, we measured minigene splicing responses of the CK8.hMbnl1V2 construct *in vitro* and observed identical splicing response as the iCAG promoter (Supplemental Figure 4B).

In agreement with the *in vitro* testing, direct IM injection showed that hNfixV2 now distinguished between WT and HSA^LR^ TA muscles (Figure 3J). Treatment with hNfixV2.MBNL1 vector showed a significant reduction in myotonic discharges and reversal of *Clasp1* exon 20 and *Clcn1* exon 7 mis-splicing, when compared to hNfixV2.eGFP-treated controls (Figure 3K-L).

### DMX^on^-controlled systemic AAV can prevent MBNL1 treatment-related toxicity

While IM delivery of MBNL1 was previously shown to improve muscle myotonia, the Chamberlain group reported severe cardiac toxicity of MBNL1 overexpression following systemic AAV-mediated delivery^21^. To evaluate the ability of DMX^on^ to prevent a treatment-related toxicity, we performed systemic delivery of DMX^on^.MBNL1 using myotropic AAV vectors (MyoAAV2A)^4^ via retro-orbital injection in WT and HSA^LR^ mice. As before, transgene expression was controlled by the CK8 promoter to drive strong expression in the heart and skeletal muscle. Importantly, the HSA^LR^ transgene is only expressed in skeletal muscle, and the ability of DMX^on^ elements to respond to endogenous MBNL1 and MBNL2 in the heart was also assessed.

Consistent with prior reports, we found that unregulated MBNL1 overexpression (MBNL1) led to severe cardiac dysfunction with reduced left ventricular ejection fraction and death after 3-4 weeks (Figure 4A-B). When DMX^on^-controlled MBNL1 vectors were delivered, hMbnl1V2 was protective while hNfixV2 was not (Figure 4A-B). Interestingly, hMbnl1V2.eGFP also led to reduced survival after 10 weeks post-injection (Figure 4A). Heart mass at tissue harvest (22-24 days post-injection for unregulated MBNL1 and hNfixV2 groups; 101-114 days for the unregulated eGFP and hMbnl1V2 groups), normalized to body weight, demonstrated cardiomegaly in unregulated MBNL1 and hNfixV2.MBNL1 mice (Figure 4C). Surprisingly, measurements in hMbnl1V2.eGFP showed a moderate reduction of ejection fraction at 4 weeks and, albeit after considerably longer exposure to the vector, enlargement of heart size at harvest (Figure 4B-C). Myotonia assessments for unregulated MBNL1 and hNfixV2.MBNL1 was limited by small sample size or short treatment duration due to cardiac toxicity, but similar to results with IM injections above, systemic delivery of hMbnl1V2.MBNL1 did not improve myotonia (Figure 4D).

**Fig 4.**
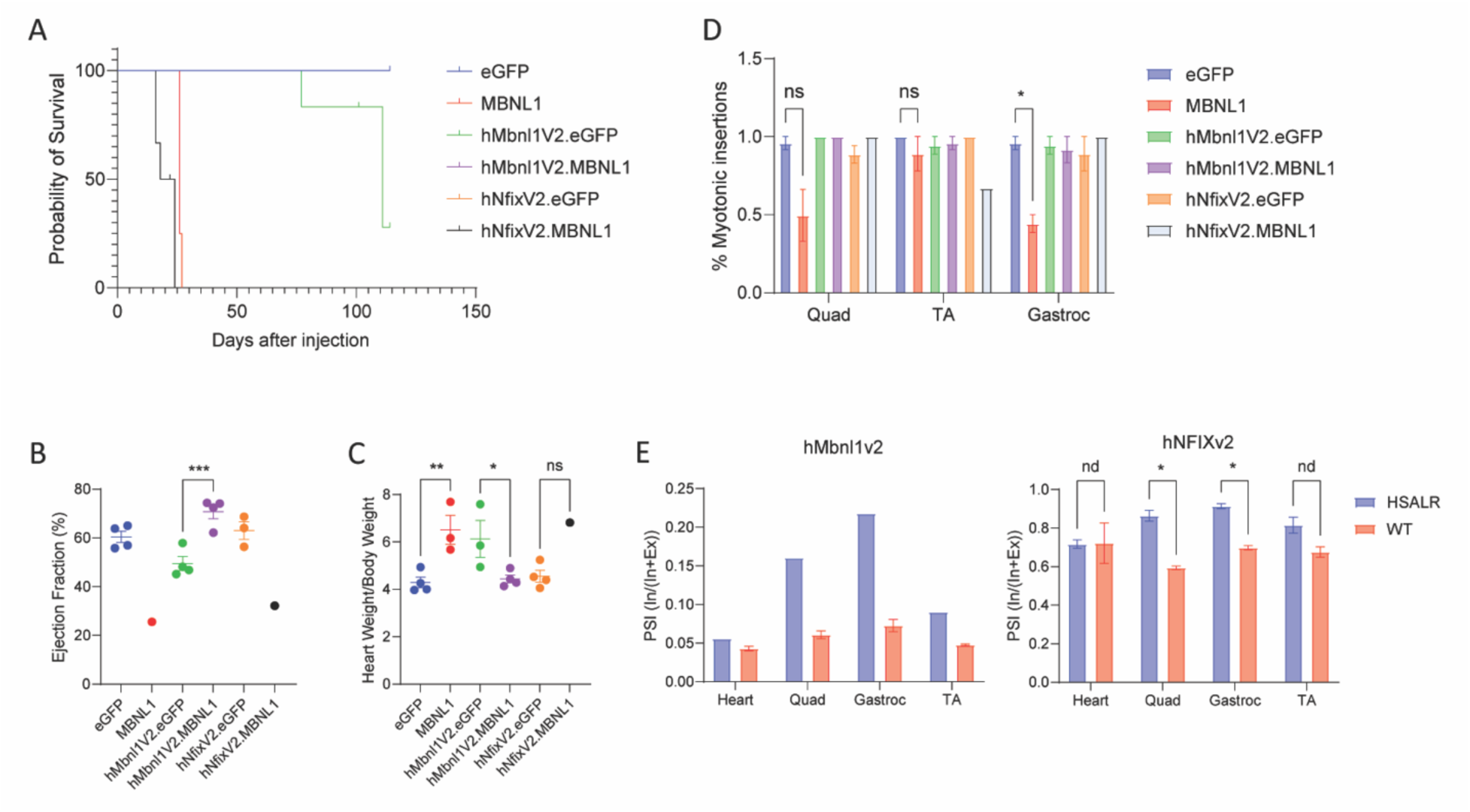
Mitigation of MBNL1 cardiac toxicity and DMX^on^ minigene responses following systemic delivery. (A) Survival curve following retro-orbital injection of vectors, as labeled [Capsid.Promoter.Minigene.Cargo] (n = 4-6 per group). (B) Quantification of ejection fraction by M-mode echocardiography 4 weeks post-injection (n = 1-4 mice/group). (C) Quantification of heart weights/body weight following harvest (n = 4/group, except hNfixV2.MBNL1 = 1). (D) Quantification of myotonia measured by needle EMG at 4 weeks post-injection (n = 1-4 mice/group). (E) Quantification of minigene PSI in the heart and three hindlimb muscles in Myo.CK8.hMbnl1V2.eGFP (left) and Myo.CK8.hNfixV2.eGFP (right) (n = 1 – 3/group).

Analysis of vector expression and DMX^on^ splicing was performed in heart, quadriceps, TA, and gastrocnemius muscles treated with eGFP-expressing vectors. Both hMbnl1V2 and hNfixV2 minigenes distinguished between WT and HSA^LR^ skeletal muscles with a PSI differential of 10-25%, depending on vector and specific hind limb muscle (Figure 4E). Interestingly, there was no significant difference in DMX^on^ transgene splicing in TA muscles between WT or HSA^LR^ mice, consistent with prior observations that the HSA^LR^ transgene expression and disease severity is variable between muscles^14^. As expected, DMX^on^ minigene splicing in the heart was similar between WT or HSA^LR^ mice and was similar to WT hind limb skeletal muscles. Vector expression of eGFP showed that the CK8 promoter drove ∼8 times higher expression in the heart compared to skeletal muscles, when normalized to *β*-actin. Consistent with use of a common promoter, expression levels among vectors were within a single logarithm with variance likely related to precision of vector titering (Supplemental Figure 5).

### DMX^on^ can distinguish between CDM and genetically corrected myotubes

To evaluate DMX^on^ response in human DM cells, we obtained an induced pluripotent stem cell (iPSC) line derived from a patient with congenital myotonic dystrophy type 1 (NH50256, NINDS Cell Repository). Isogenic control lines were generated by Cas9-mediated insertion of a polyadenylation signal upstream of the expanded CTG repeat in the DMPK 3’ untranslated region, as previously described^23,24^. This approach resulted in either insertion of the polyadenylation signal or recombination and deletion of the expanded repeat, as determined by repeat primed PCR and Cas9-targeted long read sequencing (Figure 5A and Supplemental Figure 6A). Clone 11, which showed loss of expanded repeat DNA, also demonstrated loss of nuclear foci by RNA fluorescence in-situ hybridization (FISH) using a CAG repeat probe (Supplemental Figure 6B) and was chosen as our isogenic control.

**Fig 5.**
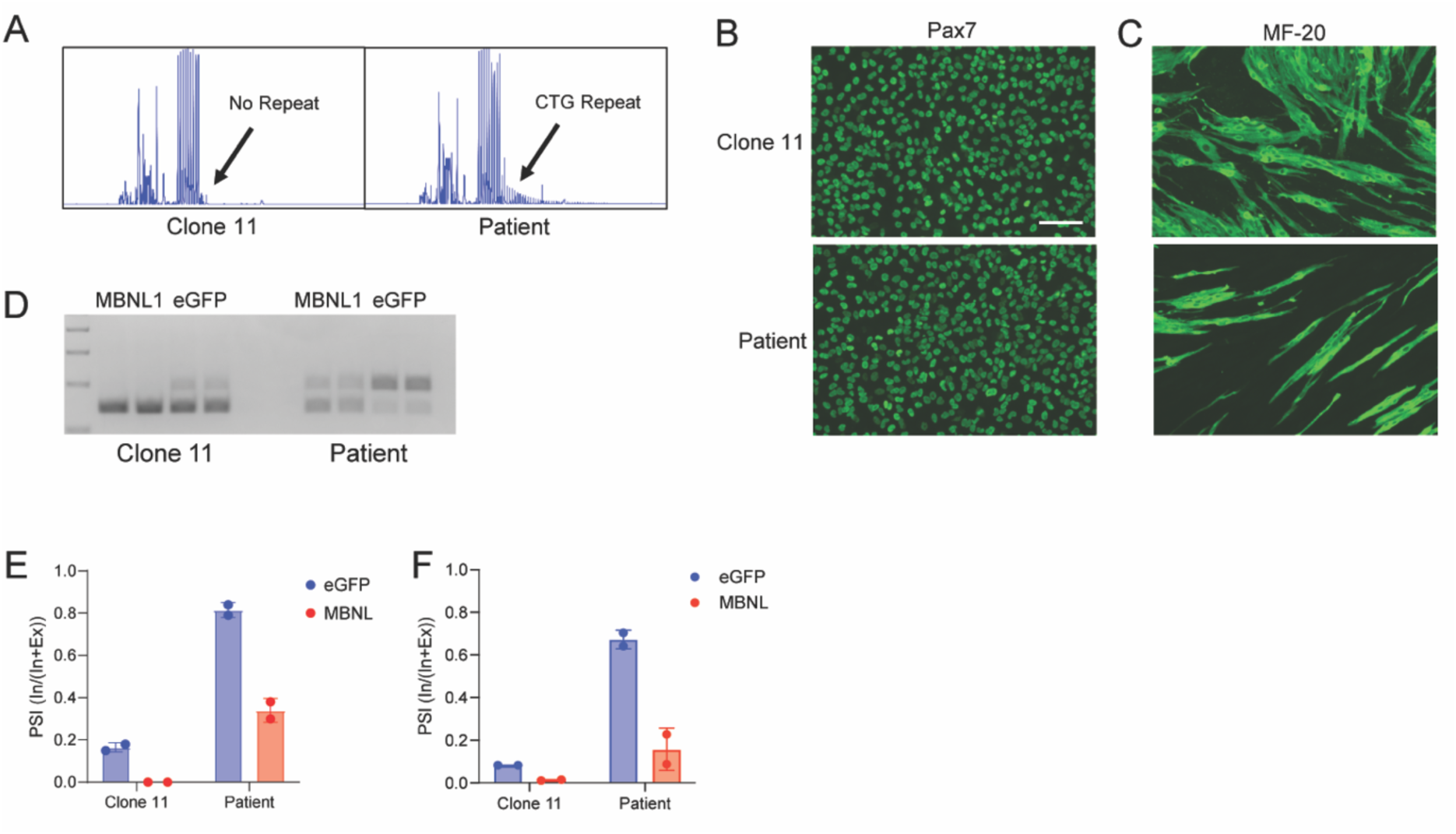
DMX^on^ response in congenital DM iPSC-derived myotubes. (A) Repeat-primed PCR demonstrating loss of repeat expansion in isogenic edited line, Clone 11. Representative images of (B) Pax7+ immunocytochemistry in myoblast cultures and (C) MF-20+ multinucleated myotubes following 10 days of differentiation. (D) Representative image of RT-PCR of hMbnl1 exon 5 inclusion in myotubes when expressing either eGFP or therapeutic MBNL1, (E) with quantification. (F) Quantification of hMbnl1 exon 5 inclusion in myotubes by ddPCR.

Clone 11 and Patient cells were programmed into myogenic precursor cells using tetracycline-inducible Pax7 overexpression^25^ before terminal differentiation into myotubes (Figure 5B). After 10 days in differentiation media, we evaluated the ability for both lines to form mature myotubes by probing with an antibody targeting myosin heavy chain (MF-20) (Figure 5C). 10 day-differentiated cultures were then treated with 1E5 vg/cell of DMX^on^ vectors under control of the hMbnl1 switch. Given the poor differentiation status of the Patient cells, we used vectors with transgenes expressed under control of the CK8 promoter to restrict expression to myotubes. Lines were incubated for an additional 7 days to allow for unpackaging and vector expression, then cells were harvested to evaluate DMX^on^ cassette response. First, to confirm the size of spliced DMX^on^ products and rule out viral gDNA contamination, we performed RT-PCR. This showed expected amplicon sizes and a clear differentiation of isogenic control cells from Patient myotubes (Figure 5D,E). We then performed ddPCR to more accurately quantify PSI (Figure 5F). The hMbnl1 switch showed strong discrimination between Patient and isogenic control cells with inclusion of the cassette exon in 64% of transcripts in Patient cells compared to only 8% in Controls. Transgene-derived MBNL1 protein expression reduced cassette exon inclusion to 23% in Patient cells and 1% in Controls, supporting negative feedback of the DMX^on^ cassette in human cells (Figure 5F).

## Acknowledgements

We thank Jean Ann Maguire and Elisa Waxman of the CHOP Human Pluripotent Stem Cell Core, Laurence Busque, and Gaige Tucker for their contributions to this work.

## Discussion

Genetic medicines have changed the therapeutic landscape for some of the most common and fatal genetic neuromuscular diseases, and viral-based gene therapy approaches have shown robust clinical efficacy^1^. A major limitation of current viral-based medicines is their inability to be re-dosed or removed once given, relying on a large therapeutic window to be safe and effective. Disease variability may greatly alter the therapeutic window from patient-to-patient in myotonic dystrophy, complicating application of a viral-based approach. To address this problem, we aimed to develop a gene therapy tuning cassette that would respond to the presence or severity of illness and adjust the gene therapy output to widen the therapeutic window.

Here, we showed that endogenous pre-mRNA alternative splicing events that are mis-regulated in DM1 can be removed from their natural context where they retain responsivity to functional MBNL1 levels or CUG repeat RNA accumulation to control output of a downstream protein. Surprisingly, the size of response in isolated DMX^on^ cassettes *in vivo* was less than that observed for their natural counterpart. This may be due to context-dependence of alternative splicing, as isolated and truncated splicing cassettes expressed as a transgene are unlikely to engage the same cadre of splicing modifiers or have the same transcriptional kinetics of their endogenous gene. Another possibility DMX^on^ cassettes generated using human sequences may be naturally less responsive than their mouse counterparts or less efficient in mouse muscle. Other factors may include the tissue targeted, the severity of disease model, the therapeutic protein being expressed, and other cargo elements, such as the promoter.

To test the functional capability of DMX^on^ elements we examined the treatment effect of MBNL1 overexpression on myotonia and RNA mis-splicing in skeletal muscle of HSA^LR^ mice and RNA mis-splicing in DM1 iPSC-derived myotubes. We chose MBNL1-41 as our therapeutic output for two primary reasons, (1) its ability to reverse skeletal muscle myotonia in the HSA^LR^ mouse model^20^ and (2) its ability to cause severe cardiac toxicity with systemic AAV delivery. Thus, when delivered at high enough titer, we could generate both a therapeutic and toxicity signal that we could attempt to modify *in vivo* with DMX^on^. We found that DMX^on^ cassettes could retain MBNL1-41 output to treat skeletal muscle myotonia and RNA mis-splicing and reduce the output of MBNL1-41 to prevent cardiac toxicity, although we did not achieve both within an individual DMX^on^ cassette using the CK8 promoter.

As well as its advantages, the decision to express MBNL1-41 as a therapeutic output is a limiting factor in this study for multiple reasons. First, the overexpression of MBNL1-41 can bind to and directly feed back onto DMX^on^. This may more strongly inhibit DMX^on^ activity than an alternative treatment that indirectly affects DMX^on^ by releasing endogenous MBNL1.Second, although expanded repeat RNAs can self-aggregate and form strong condensates *in vitro*^26^, evidence *in vivo* and from experiments with MBNL knockdown or overexpression suggests that the amount of total toxic RNA aggregates is MBNL-dependent^20,27^. This may explain the observation that the therapeutic effect of MBNL1-41 overexpression wanes after 6 months^20^, but may also be due to enhanced muscle regeneration, as previously suggested.

Another limitation may be the HSA^LR^ model as it is not an ideal model to validate DMX^on^ responses. As previously reported and observed in our data, hind limb muscles are differentially affected by the HSA^LR^ transgene, for example, quadriceps are more affected than tibialis anterior^28^, a pattern opposite that observed in human DM1. Additionally, despite a ∼5 times greater CUG repeat RNA load than that observed in patient muscle samples^29^, HSA^LR^ mice demonstrate moderate disease phenotypes^14^, suggesting unique biophysical characteristics and condensate formation involved with longer repeat expansions. Further, it is unclear how a mildly affected or clinically unaffected heart in a patient with DM1 would relate to unaffected heart in an HSA^LR^ mouse, as recent data suggests that MBNL1 overexpression is beneficial in murine cardiac models of DM1, not detrimental (Correspondence with Dr. Thomas Cooper).

Last, our overall approach to develop DMX^on^ was by using a candidate gene approach of known MBNL-dependent splicing events. An alternative approach would be to screen for synthetic alternative splicing cassettes *in vivo* using an iterative library of potential variants. This approach may identify a more responsive DMX^on^ cassette, although the variability between tissues or in response to a chosen treatment modality is likely to remain.

One aspiration of our approach that was not assessed in the current study is the ability of DMX^on^ to control gene therapy with a ubiquitously expressed promoter and more broadly tropic capsid. MBNL proteins are broadly expressed throughout the body, but are significantly upregulated during development most clearly in brain, heart, and skeletal muscle^30^. Expression data from the Human Protein Atlas suggests that either MBNL1 or MBNL2 is also expressed in skin/connective tissue, endocrine tissues, eye, liver, and the digestive tract. It is likely that empiric testing would be required to better understand the responsiveness of our MBNL-dependent splicing event in these tissues.

Taken together, our data support that AAV cargo elements can respond to and post-transcriptionally tune output of a delivered gene therapy, dependent on the disease-state of the target tissue. These elements may have limited dynamic range, and while modifiable to adjust their response, are subject to tissue-specific and treatment-specific effects which would need to be tested empirically.

## Methods

### Plasmids and AAV vectors

Genomic sequence from *LDB3*, *MBNL1*, and *NFIX* were synthesized by gBlock using hg38 as a reference genome. Recombinant AAV serotype AAV2/9 or MyoAAV2A were generated at the Children’s Hospital of Philadelphia Research Vector Core and resuspended in Diluent buffer. Vector titers were determined by ddPCR using primers targeting eGFP or MBNL1 transgenes. For MBNL1 therapeutic overexpression vectors, sequence was amplified from Addgene plasmid # 96906; http://n2t.net/addgene:96906 ; RRID:Addgene_96906; a gift from Thomas Cooper.

### Induced pluripotent stem cell culture

The induced pluripotent stem cell (iPSC) line NH50256 was obtained from the NINDS repository (stemcells.nindsgenetics.org). This line was generated from a type 1 congenital myotonic dystrophy patient with ∼1150 CTG repeats. Prior to editing, the line underwent testing for Sendai viral vector clearance, mycoplasma, karyotyping, trilineage potential, and markers of stemness (SSEA and Tra) with the CHOP Stem Cell Core. To make isogenic control cells, Cas9-mediated targeted insertion of a polyadenylation signal upstream of the expanded CTG repeat was performed, as previously described^31^. Multiple clones were generated from this editing, the line chosen for these experiments (Clone 11) resulted in deletion of the expanded repeat, confirmed by long-read sequencing. Lines were maintained in feeder-free conditions with mTeSR1 on Matrigel and passaged by clump method.

### Myoblast and myotube differentiation

Myoblasts/myotubes were generated by a combined inhibitor and PAX7 transgene overexpression method, as previously described^32^. In brief, iPSCs were co-transduced with FUGW-rtTA and pSAM2-iPAX7-IRES-tomato lentiviral vectors (kind gift of Rita Perlingeiro, PhD). Embryoid bodies (EBs) were formed by incubating cells on orbital shaker with ROCK inhibitor (Y-27632) for 2 days. EBs were then treated with CHIR99021 (GSK3*β* inhibitor) in myogenic media (previously described) for 2 days, then BMP (LDN193189) and TGF*β* (SB431542) inhibitors were added. One day later doxycycline was added to induce PAX7 expression for 2 additional days. EBs were then plated on gelatin-coated plates and expanded in myogenic media supplemented with basic fibroblast growth factor (bFGF) for 4 days, then sorted for TdTomato expression to enrich for myogenic progenitors. These progenitors were then frozen prior to performing terminal differentiation for experiments. For terminal differentiation, progenitors at passage 2+ were plated at 100-120,000 cells per well of a 24-well plate and grown to high confluency for 3 days. Media was then changed to differentiation media (as previously described) and monitored for 5-10 days for formation of myotubes.

### Mouse models and viral injections

All mice were bred and housed within an accredited AALAC facility. All procedures were approved by the institutional IACUC committee prior to experiments. Wild-type animals were FVB/NJ strain and HSA^LR^ mice were the FVB/N-Tg(HSA*LR)20bCath/J transgenic line (Jackson Laboratory). HSA^LR^ mice are transgenic for the human ACTA1 gene and proximal promoter with ∼220 CTG repeats inserted into the 3’ untranslated region. The HSA^LR^ line was maintained as homozygous breeders, CTG repeat lengths were monitored in litters prior to selection of breeders, and periodic outcrossing back to FVB/NJ was performed.

Direct intramuscular (IM) injections were performed in tibialis anterior muscles in mice. In brief, mice were anesthetized with isoflurane, distal hind limbs were shaved, and 5E9 viral genomes (vg) was brought to 20 µl in Diluent buffer and was injected into the TA with a 30-gauge insulin syringe.

Systemic delivery was performed by retro-orbital injection. In brief, mice were anesthetized with isoflurane, 0.5% proparacaine anesthetic eye drops were delivered to the right eye and allowed to take effect for 2-5 minutes, a 30-gauge insulin syringe was then passed into the retro-orbital sinus, and 2E13 vg/kg of viral vector in Diluent buffer to no more than 200 µl was injected.

### Immortalized cell culture

HEK293 cells were previously engineered to express tetracycline-inducible HA-MBNL1 protein^17^. This line was used to test *in vitro* splicing and protein output of DMX^on^ minigenes. Cell cultures were maintained with DMEM (Gibco), 10% Tet-System Approved FBS, 1% Penicillin/Streptomycin. For both RNA and protein experiments, 200,000 cells were plated in each well of a 24-well plate. The following day, cells were transitioned to doxycycline-containing media and transfected 4 hours later with 250 ng plasmid/well using Lipofectamine 2000, per manufacturer’s protocol. Doxycycline-containing media was replaced every 24 hours for experiments exceeding 24 hours.

### RNA isolation and Reverse Transcription

For *in vitro* splicing assays in HA-MBNL1 HEK293 cells, RNA was isolated 24 hours after plasmid transfection by Trizol, DNAse treated using DNA-free kit (Ambion), and 1 ug of RNA was reverse transcribed using a High Capacity cDNA Reverse Transcription kit (Applied Biosystems), per manufacturer’s protocol. Individual muscle or heart samples were homogenized using stainless steel beads in the TissueLyser LT (Qiagen) in Trizol, then similarly treated with DNase, before 2 ug of RNA was reverse transcribed. iPSC-derived myotube RNA was isolated and DNase treated using the Zymo Quick RNA Miniprep kit. Reverse transcription used the High Capacity cDNA Reverse transcription kit with variable input, depending on recovered concentration.

### Digital Droplet PCR (ddPCR) analysis of minigene splicing

PCR primers were designed to amplify the region of the DMX^on^ switches (CK8_Mbnl_ddPCR_For: AGCGAATTAAACTCGAGACCA or CAG_ddPCR_For: TGT GCT GTC TCA TCA TTT TGG; Mbnl_ddPCR_Rev: GGGAAGTACAGCTTGAGGAAT; CK8_Nfix_ddPCR_For: AGCCAGCCAGCGAATTAAA or or CAG_ddPCR_For: TGT GCT GTC TCA TCA TTT TGG; Nfix_ddPCR_Rev: GGAAGTGCAGGGCTGATG). HEX-labeled probes were designed to bind to the alternate cassette exon (Mbnl_ddPCR_Cassette: /5HEX/AATCACTGA/ZEN/AGCCACCATGCTCGA/3IABkFQ/; NFIX_ddPCR_Cassette: /5HEX/AGTCAGGAA/ZEN/AGCTGGACTTCTGCA/3IABkFQ/) and FAM-labeled probes to common sequence in flanking exons (Mbnl_ddPCR_Common: /56-FAM/CAACCAGGC/ZEN/TGCAGC TGCA/3IABkFQ/; NFIX_ddPCR_Common: /56-FAM/CGTCCACTT/ZEN/CGTTGGGCCAC/3IABkFQ/). All oligos were ordered from Integrated DNA Technologies. cDNA samples were diluted up to 50x to keep measurements within the linear quantification range, and mixed with Supermix (Bio-Rad), primers, and probes. Droplets were read using the QX200 ddPCR system (Bio-Rad).

### Semi-quantitative RT-PCR of alternative splicing

PCR primers designed across mouse *Serca1* exon 21-23 (For: CTCATGGTCCTCAAGATCT CAC, Rev: GGGTCAGTGCCTCAGCTTTG), *Clcn1* exon 5-8 (For: TGAAGGAATACCTCACAC TCAAGG, Rev: CACGGAACACAAAGGCACTG), and *Clasp1* exon 19-21 (For: GCCAGTGCCA AATCCAAAG, Rev: GCTGAGACTGTGAAACCACT) were used to amplify cDNA generated from experimental animals using PrimeSTAR GXL DNA polymerase (Takara Bio). Products were visualized using 2.5% agarose gel electrophoresis stained with ethidium bromide. Band intensity was measured with Image J.

### Quantitative RT-PCR (qPCR)

Vector expression levels were measured using a CFX384 Real Time System (Bio-Rad) using TaqMan Universal Master Mix II, no UNG (Applied Biosystems) with primer/probe set directed against eGFP (For: GAACCGCATCGAGCTGAA, Rev: TGCTTGTCGGCCATGATATAG, Probe: /56-FAM/ATCGACTTC/ZEN/AAGGAGGACGGCAAC/3IABkFQ/), normalized to mouse *β*-actin (Mm00607939_s1).

### Western Blot

Cell lysates were harvested by cell scraping in cold RIPA buffer (150 mM NaCl, 1% Igepal, 0.5% NaDOC, 0.1% SDS, 50 mM Tris pH 8.0) supplemented with 1X cOmplete protease inihibitors (Roche). Protein concentrations were measured using a Bio-Rad RC DC assay and 20 µg of protein was then mixed with XT Sample Buffer (Bio-Rad) and XT Reducing Agent (Bio-Rad) and heated at 95 C for 5 minutes. SDS-PAGE was then performed on Criterion XT 10% Bis-Tris gels (Bio-Rad), transfer to PVDF membrane was performed overnight at 4C in Towbin buffer. Membranes were blocked with 5% non-fat milk in TBST before primary incubation with either 1:5000 anti-MBNL1 (A2764), kind gift from Charles Thornton) or 1:5000 anti-eGFP antibodies (abcam, ab290), both dilute in block solution. Secondary HRP-conjugated antibodies were dilute in 5% milk and incubated at 1:10,000 for 1 hour at RT and exposed by Amersham ECL detection kit (Cytiva, RPN2105).

### Needle Electromyography

A Natus Nicolet EDX machine was used to perform needle EMG to assess myotonia in HSA^LR^ mice. Mice were anesthetized by isoflurane and maintained on a heating pad. Natus Teca Elite concentric needle electrode (length: 25mm, diameter: 0.3 mm) was then inserted into hind limb muscles to assess for myotonic discharges. Six total insertions were performed and video recorded per muscle. The percentage of insertions leading to a myotonic discharge was tallied.

### Echocardiography

Mice were examined under isoflurane anesthesia, maintained on a warmed platform with electrocardiogram monitoring and heart rates above 400 bpm. M-mode echocardiographic recordings were performed on short-axis videos at the mid-papillary level as well as short- and long-axis 2D videos. These were used to measure left ventricular wall thickness and internal diameter, then calculate ejection fraction, cardiac output and mass. All measurements and analysis of echocardiography were performed by a skilled and blinded echo technician.

