## Supplemental Figures for "Co-opting MBNL-dependent alternative splicing cassette exons to control gene therapy in myotonic dystrophy"

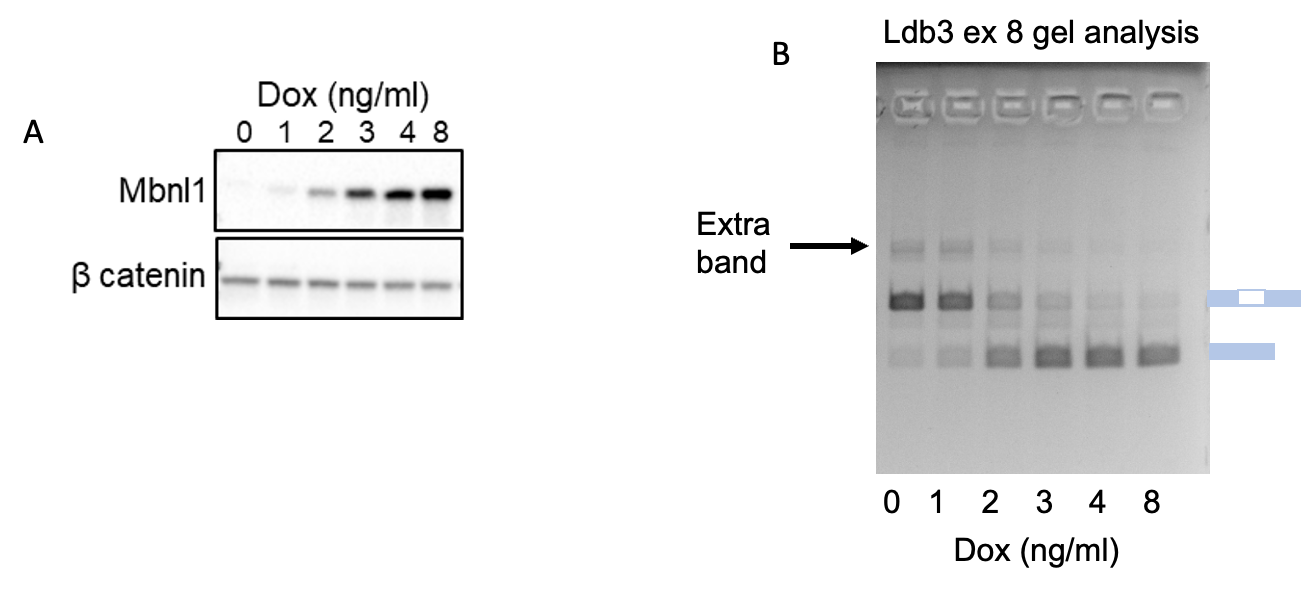

**Supplemental Figure 1. Doxycycline-inducible MBNL1 HEK293 cells and Ldb3 abnormal splicing.** (A) Representative Western blot of Mbnl1 protein expression in tetracycline-inducible HA-MBNL1 HEK 293 cells across a range of doxycycline concentrations. (B) Representative agarose gel of RT-PCR to measure alternative splicing of Ldb3 exon 8, arrow demarcated larger product.

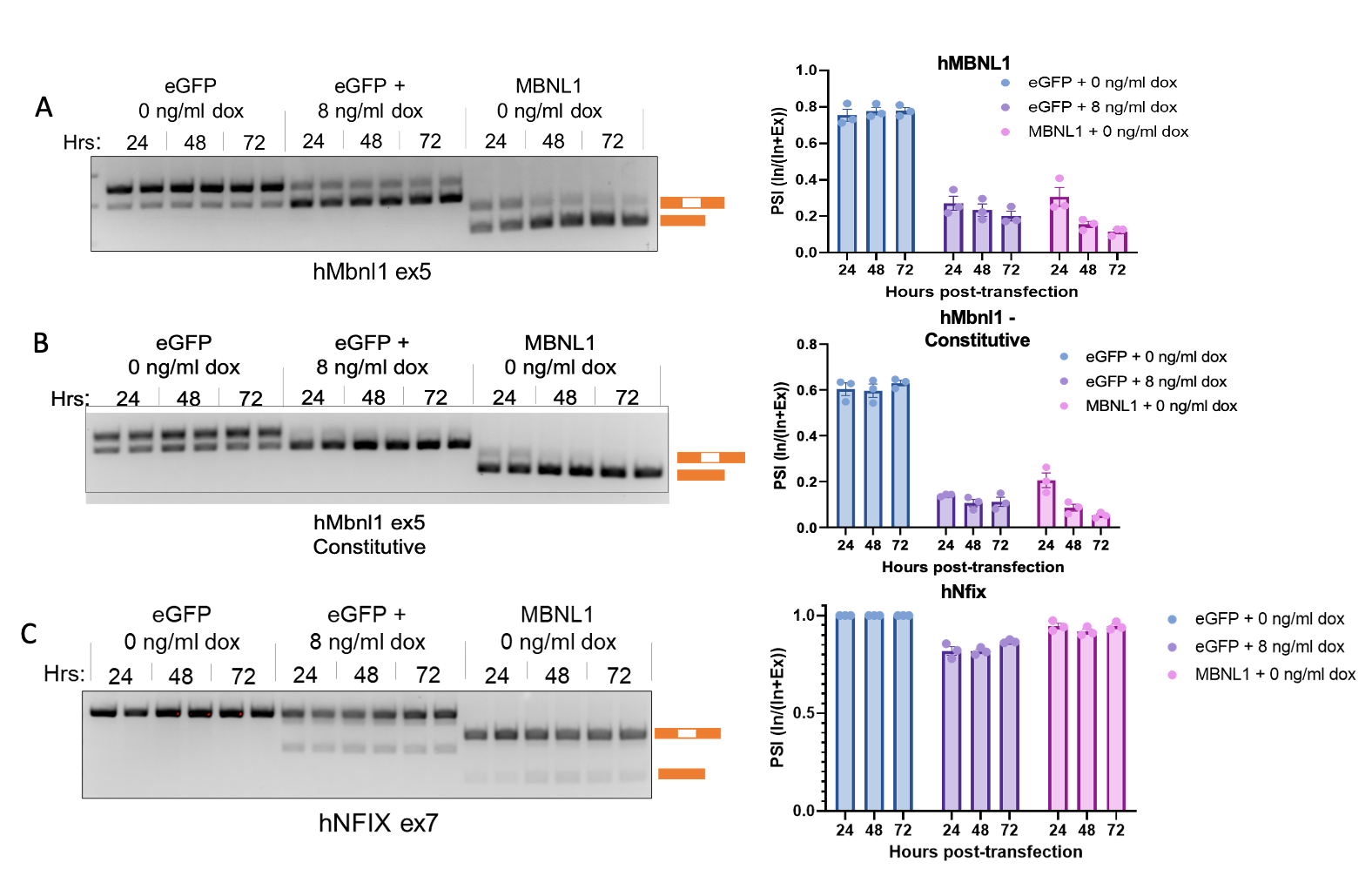

**Supplemental Figure 2.** **DMX^on^ self-regulation to MBNL1 overexpression.** Representative RT-PCR agarose gels of (A) hMbnl1 exon 5, (B) hMbnl1 exon 5 constitutive, and (C) hNfix exon 7 splicing over a 72-hour time course in response to MBNL1 protein driven in HA-MBNL1 tet-inducible cells or from self-expression of MBNL1.

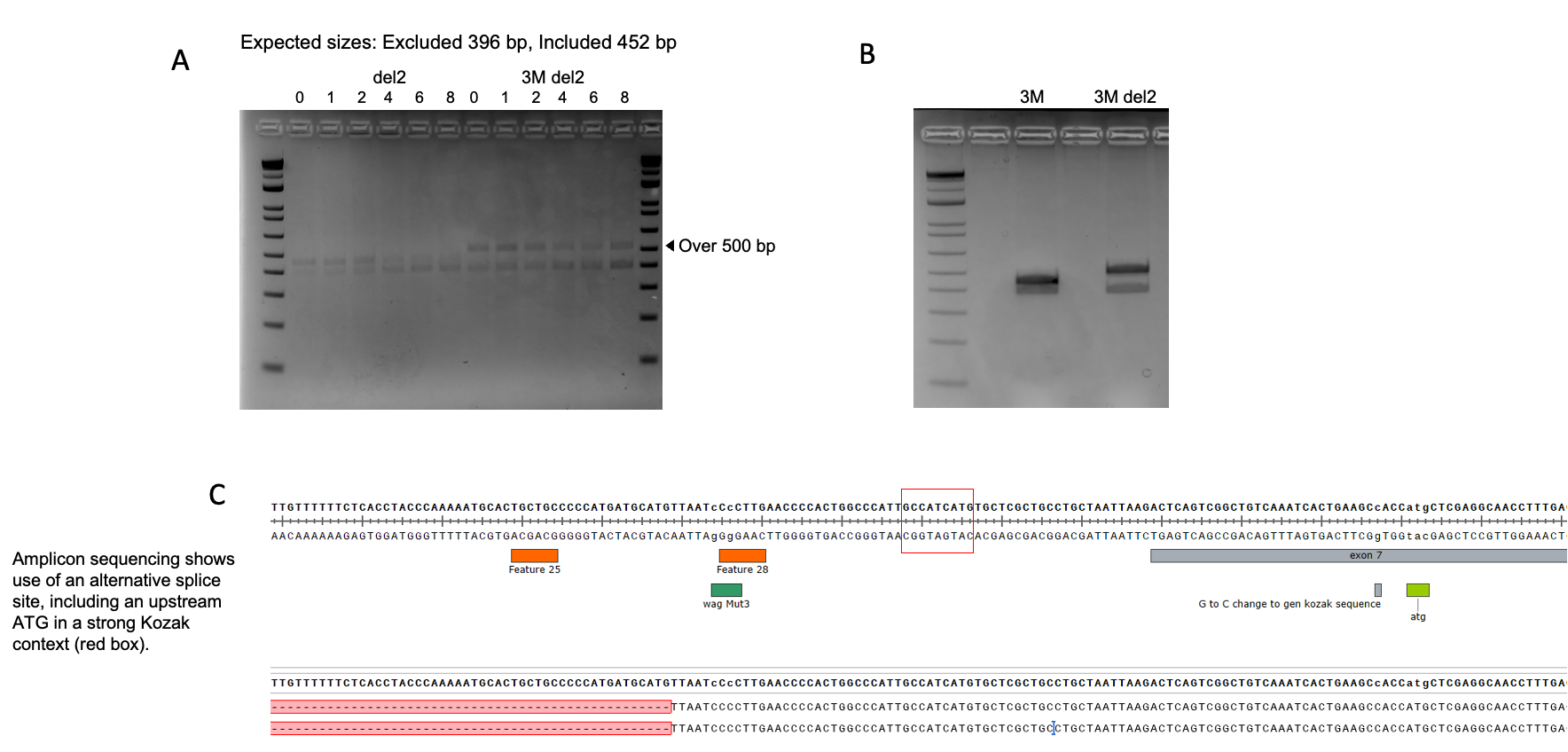

**Supplemental Figure 3. Aberrant use of upstream alternative splice site in combined hMbnl1 variant.** (A) Representative RT-PCR gel of hMbnl1 del2 and 3M+del2 across a range of doxycycline. (B) Same as (A), but without doxycycline in order to isolate the larger amplicon for sequencing. (C) Sequence of intron 6 – exon 7 junction with Sanger sequencing below showing inclusion of upstream sequence and strong Kozak sequence and start codon (red box).

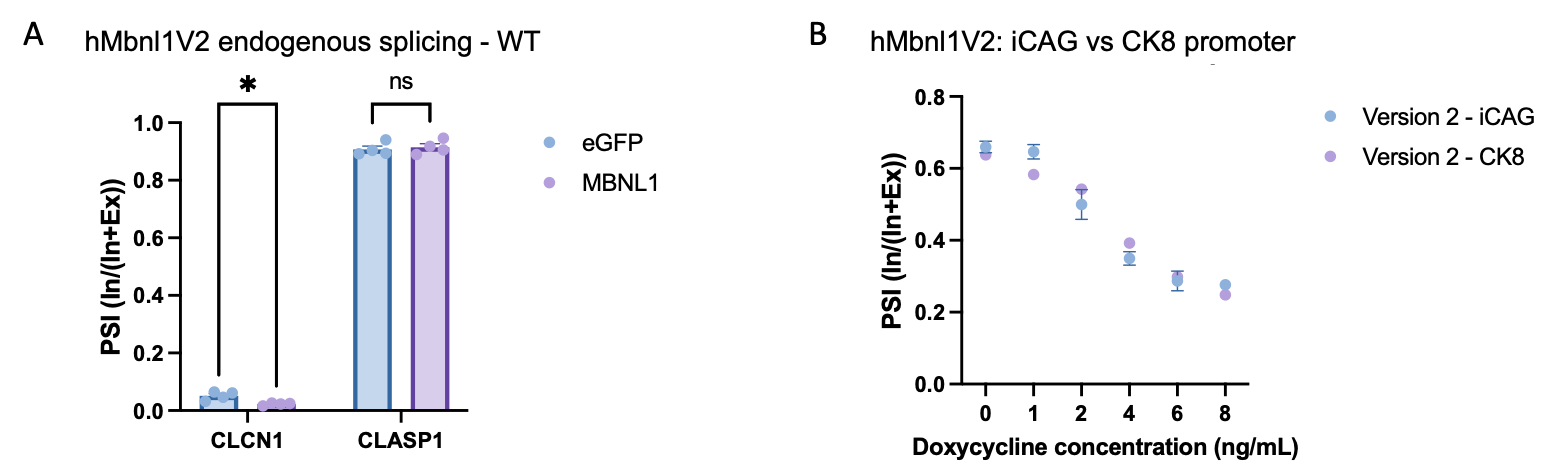

**Supplemental Figure 4. hMbnl1V2 minor impact of endogenous splicing in WT mice and reporter splicing is not reliant on promoter.** (A) Quantification of endogenous splicing of mouse CLCN1 exon 7 and CLASP1 exon 20 in eGFP or MBNL1 treated TA muscles in WT mice (n = 4 / group). (B) Quantification of PSI by ddPCR across a range of doxycycline in tetracycline-inducible HA-MBNL1 HEK 293 cells when driven by either iCAG or CK8 promoter.

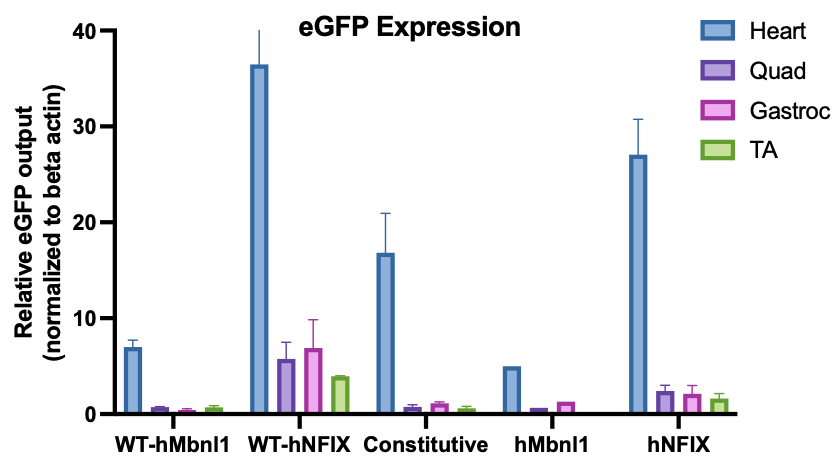

**Supplemental Figure 5. eGFP expression is similar between vectors when delivered systemically.** Quantification of eGFP expression in various tissues following systemic delivery of constitutive or DMX^on^ regulated vectors in WT or HSALR mice.

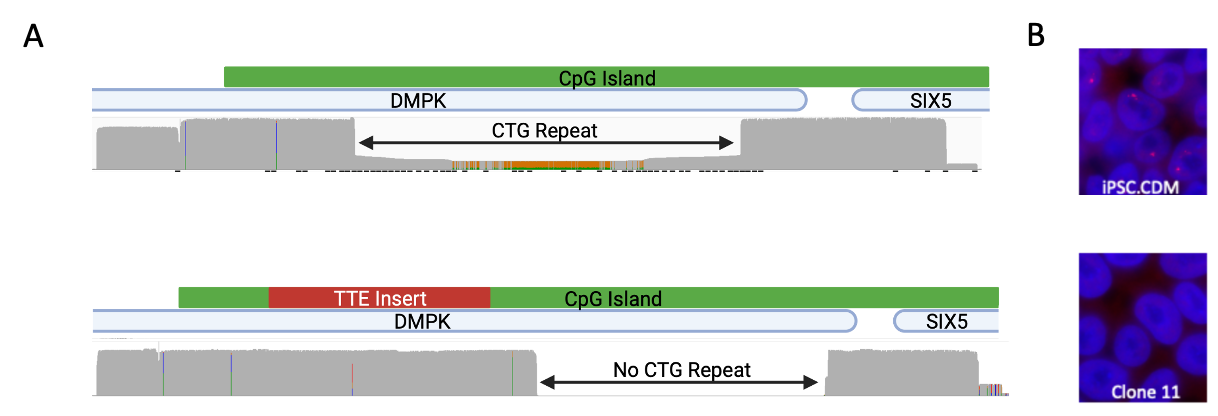

**Supplemental Figure 6. Insertion of polyadenylation signal by Cas9-mediated targeting leads to CTG repeat deletion and loss of RNA foci in CDM iPSCs.** (A) Coverage maps of long-read sequencing of DMPK repeat region in congenital DM1 patient iPSC (above) and edited clone 11 (below). (B) Representative images of RNA fluorescence in situ hybridization using a Cy3-conjugated (CAG)5 probe (PNA Bio) in CDM iPSC or edited clone 11 cells.
